# CAT1/SLC7A1 acts as a cellular receptor for bovine leukemia virus infection

**DOI:** 10.1101/666958

**Authors:** Lanlan Bai, Hirotaka Sato, Yoshinao Kubo, Satoshi Wada, Yoko Aida

## Abstract

Bovine leukemia virus (BLV) is the causative agent of enzootic bovine leukosis, the most common neoplastic disease of cattle, which is closely related to human T-cell leukemia viruses. BLV has spread worldwide and causes a serious problem for the cattle industry. The cellular receptor specifically binds with viral envelope glycoprotein (Env) and this attachment mediates cell fusion to lead virus entry. BLV Env reportedly binds to cationic amino acid transporter 1 (CAT1)/SLC7A1, but whether the CAT1/SLC7A1 is an actual receptor for BLV remains unknown. Here, we showed that CAT1 functioned as an infection receptor, interacting with BLV particles. Cells expressing undetectable CAT1 levels were resistant to BLV infection but became highly susceptible upon CAT1 overexpression. CAT1 exhibited specific binding to BLV particles on the cell surface and co-localized with the Env in endomembrane compartments and membrane. Knockdown of CAT1 in permissive cells significantly reduced binding to BLV particles and BLV infection. In addition, bovine serum with neutralizing activity from a BLV-infected cattle inhibited BLV particles. Expression of CAT1 from various species demonstrated no species-specificity for BLV infection, implicating CAT1 as a functional BLV receptor responsible for its broad host range. These findings provide insights for BLV infection and for developing new strategies for treating BLV and preventing its spread.

**Author Summary:** Bovine leukemia virus (BLV), which can infect a variety of animal species and induce lymphoma in cattle, is a member of the family *Retroviridae*. BLV induces huge economic losses by not only lymphoma but also subclinical forms of the disease. In addition, BLV is frequently used as an animal model of human T-cell leukemia virus (HTLV), as BLV has many similar characteristics to HTLV. Thus, understanding BLV pathogenesis contribute to resolve not only BLV-but also HTLV-induced problems. Retroviral envelope glycoprotein (Env) is specifically recognized by the cellular receptor at cell surface, which induces a conformational changes between viral and cell membrane to entry. Thus, the elucidation of cellular receptor for BLV infection is very important for virus entry. However, the BLV receptor has not been identified yet. In the current study, we found that BLV Env protein binds to cationic amino acid transporter 1 (CAT1)/SLC7A1 at cell surface, artificial expression of CAT1 in CAT1-negative cells confers the cells susceptible to BLV infection, and CAT1-silencing significantly reduces BLV infection, concluding that CAT1 is the BLV receptor. These findings will have far reaching great advantages of insights in the retrovirus study.

## Introduction

Bovine leukemia virus (BLV) is the causative agent of enzootic bovine leukosis, a B-cell leukemia/lymphoma [1–4] that has spread worldwide and causes serious problems for the cattle industry [5]. The natural hosts of BLV are domestic cattle and water buffaloes [1] but BLV can also be experimentally transmitted to several other animal species, such as sheep [6, 7], goats [8], pigs [9], rats [10], rabbits [11], and chickens [12]. However, BLV induces leukemia/lymphoma only in cattle and sheep [1, 4]. Importantly, BLV has been detected in the breast epithelium of humans [13–16]. In addition to the many studies showing that the BLV host range is broad, BLV can also successfully infect a variety of cells *in vitro* [17].

Like other retroviruses, the BLV genome comprises *gag, pro, pol*, and *env*, as well as regulatory genes *tax* and *rex* and accessory genes *R3* and *G4* [1, 4]. The Gag protein encoded by *gag* plays important structural roles in the assembly of virions at the plasma membrane and in genome packaging. BLV *env* encodes precursor Pr72 envelope glycoprotein (Env), which is glycosylated in the rough endoplasmic reticulum and Golgi apparatus [18] and is processed into two mature proteins, the surface glycoprotein (SU) subunit gp51 and the transmembrane (TM) subunit gp30 [19, 20]. The gp51 and gp30 proteins form a stable complex through disulfide bonds [20] and are incorporated into budding viral particles [21]. Gp51 contains three conformational epitopes (F, G, and H) in its N-terminus that are recognized by neutralizing antibody [4, 22]. The C-terminus of gp51 also contains four linear epitopes (A, B, D, and E) [23]. In addition, there is evidence that the N-terminal half of mature gp51 plays important roles in virus infectivity and syncytium formation, suggesting that it probably contains the receptor-binding domain (RBD) [20]. However, the actual cellular receptor for BLV infection has not been identified.

The *env* genes of BLV and its close relative human T-cell leukemia virus type 1 (HTLV-1) are highly conserved [1, 4]. In particular, their Env proteins show 36% identity in their amino acid sequences [4]. Entry of these retroviruses into target cells is initiated by interaction between Env and host cellular receptors [24, 25]. Glucose transporter 1 (GLUT1) [26], neuropilin-1 (NRP-1) [27], and heparan sulfate proteoglycans (HSPG) [28] have been determined to be cellular receptors for HTLV attachment and/or infection. When NRP-1, GLUT1, and the SU subunit of HTLV-1 are co-expressed in cells, they are able to form a stable tripartite complex [27]. The initial attachment and concentration of HTLV-1 virions at the cell surface [29] are induced via interaction of HSPG with the C-terminal domain of the SU subunit of HTLV-1 [30]. HSPGs also directly bind with NRP-1, which forms a complex with HTLV-1 Env to induce conformational changes in the SU subunit allowing residues D106 and Y114 of SU to bind with GLUT1 [26, 30]. This multi-receptor usage is required for HTLV-1 entry into target cells [30, 31].

GLUT1 is composed of 12 hydrophobic TM domains, six extracellular loops (ECL), and seven intracellular domains [32]. Similar to GLUT1, cationic amino acid transporter 1 (CAT1)/SLC7A1 has 14 potential membrane-spanning domains and has been identified as a membrane receptor on mouse cells for ecotropic murine leukemia viruses (eMuLVs) [33]. CAT1 is a 622-amino acid sequences of protein that is extremely hydrophobic and implicated in sodium-independent transport of arginine, lysine, and histidine [33–35]. Two distinct motifs (YGE and HE) in the third extracellular loop of CAT1 are known to bind with the N-terminus of the eMuLV SU subunit for viral entry [36, 37] into mouse and rat cells [38], indicating that the third extracellular loop in CAT1 is a determinant for eMuLV infection. CAT1 of human cells does not provide susceptibility to eMuLV infection; however, expressing mouse CAT1 in human cells can result in acquired susceptibility [39]. Similar to human cells, hamster cells are completely resistant to eMuLV infection [38] and CAT1 proteins from other animals are also resistant to eMuLV infection, indicating CAT1 may provide the species-specificity for eMuLV infection.

At the 29th International Workshop on Retroviral Pathogenesis (2017), Dr. Marc Sitbon presented BLV Env binds to CAT1; however, it is not clear whether CAT1 actually serves as a cellular receptor for BLV. In the current study, we detected CAT1 protein in various cells and its correlation with susceptibility to BLV infection. Next, using bovine CAT1/SLC7A1 expression plasmid, we investigated whether exogenous expression of CAT1 induces BLV infection in CAT1-negative, BLV-resistant cells. In addition, we determined the CAT1 and BLV particles binding and co-localization with the Env in the cells. Furthermore, we analyzed the impacts of CAT1 knockdown on BLV cell-free and cell-to-cell infection and BLV particles binding. Finally, we clarified the species-specific susceptibility of CAT1 for BLV infection.

## Results

### Detection of CAT1 protein in various cells from multiple animal species and its correlation with susceptibility to BLV infection

To verify whether CAT1 is responsible for the broad host range of BLV *in vitro*, CAT1 expression in eight cell lines derived from six animal species was first measured using western blot analysis with an anti-CAT1 antibody. As shown in Fig. 1, the human cervical HeLa cells, human embryonic kidney 293T cells, feline CC81 cells, calf kidney CKT-1 cells, bovine lymphoid KU-1 cells, permanently BLV-infected fetal lamb kidney (FLK-BLV) cells, and porcine kidney PK15 cells each expressed CAT1. The Chinese hamster ovary CHO-K1 cells were the only cell line that did not express detectable levels of CAT1 protein.

**Fig. 1.**
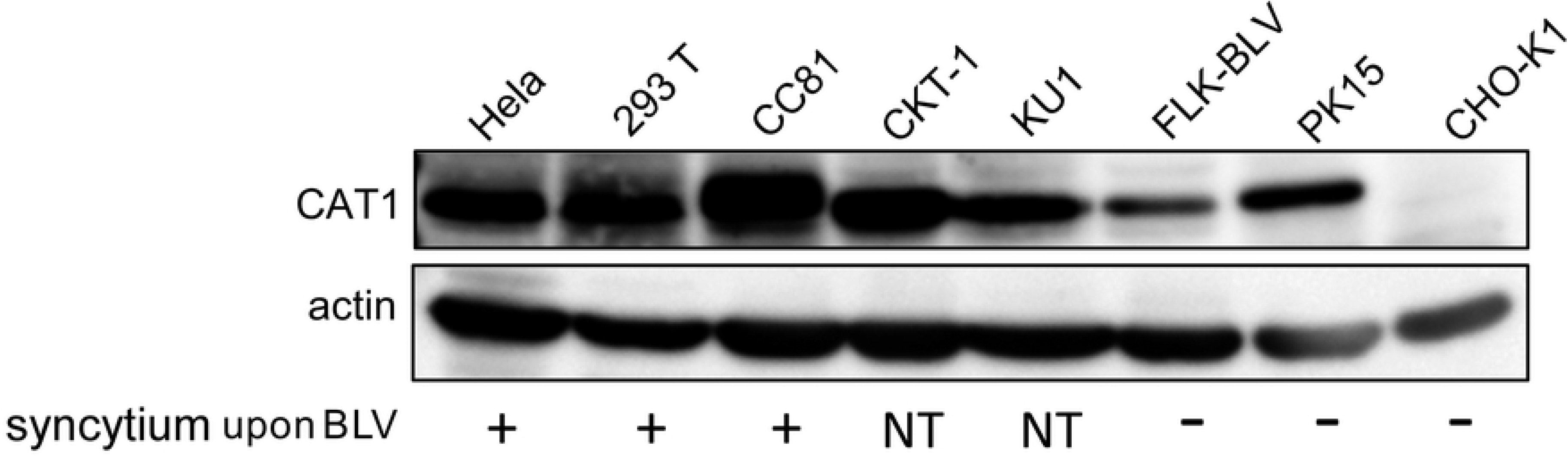
Detection of CAT1 expression in eight different species-based cell lines and its correlation with susceptibility to infection by bovine leukemia virus (BLV). Cell lysates were subjected to western blot analysis using anti-CAT1 and anti-β-actin antibodies to detect CAT1 expression in each cell line (upper panel). The molecular weights of CAT1 and actin are indicated. The cell lines were infected by co-culturing with BLV-infected cells or by transfection of the infectious molecular clone pBLV-IF2 (lower). The results of syncytium formation for each cell line following BLV infection are shown as positive (+), negative (−), and not tested (NT).

As BLV-infected cells fuse with neighboring cells thereby forming syncytia in culture, BLV infection is typically monitored *in vitro* using syncytium formation. To evaluate a correlation between CAT1 expression and susceptibility to BLV infection, we inoculated each cell line with BLV by co-culturing them with BLV-infected cells or by transfection with the infectious molecular clone pBLV-IF2 (Fig. 1). Inoculation with BLV induced syncytia in HeLa, 293T, and CC81 cells, which expressed CAT1 at high levels, but not in PK15 cells, which expressed CAT1 at low levels. No syncytia were detected in FLK-BLV cells since the CAT1 was already occupied by the BLV Env protein. In contrast, CHO-K1 cells, which did not express detectable levels of CAT1, failed to form syncytia upon inoculation with BLV. These results indicated that CAT1 expression correlated with cellular susceptibility to BLV infection.

### CAT1/SLC7A1 expression induced BLV infection

To investigate the role of CAT1 as a receptor for BLV, we constructed an EGFP-tagged bovine CAT1 expression plasmid (bCAT1/pEGFP-N1) and CAT1 expression was visualized. A single band of endogenous CAT1 with the expected molecular size of 68 kDa was detected in positive control CC81 cells using an anti-CAT1 antibody, but no specific bands were detected in non-transfected or pEGFP-N1-transfected CHO-K1 cells. In bCAT1/pEGFP-N1-transfected CHO-K1 cells, a protein band of the expected molecular size of 95.5 kDa was believed to be a glycosylated EGFP-tagged bovine CAT1 protein with several additional bands of lower molecular sizes also detected by western blotting using an anti-CAT1 antibody (Fig. 2A, upper panel) and anti-EGFP antibody (Fig. 2A, middle panel). The lower molecular size bands were thought to be glycosylated variants.

**Fig. 2.**
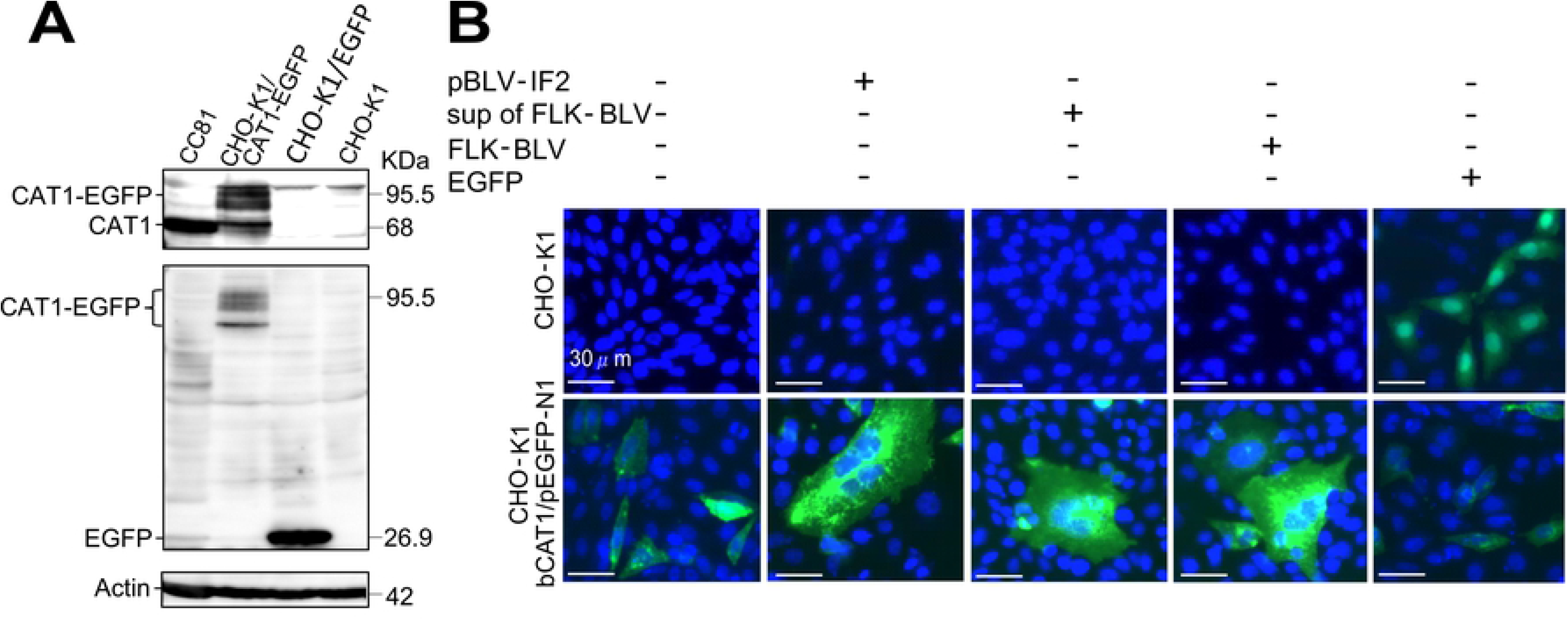
CAT1/SLC7A1 protein acted as a cellular receptor and was required for bovine leukemia virus (BLV) infection. (A) Confirmation of the absence of endogenous CAT1 expression in CHO-K1 cells. CHO-K1 cells were transfected with and without bCAT1/pEGFP-N1 or with pEGFP-N1 as a negative control. The cell lysates were prepared 48 h post transfection and then subjected to western blot analysis using anti-CAT1 (upper panel), anti-EGFP (middle panel), and anti-β-actin (lower panel) antibodies. CC81-GREMG cell lysates were used as a positive control. The positions and molecular weights of CAT1-EGFP, CAT1, EGFP, and actin are indicated. (B) Bovine CAT1/SLC7A1 expression associated with BLV cell-free and cell-to-cell infection. CHO-K1 cells were transfected with bCAT1/pEGFP-N1 expression plasmid with or without the BLV infectious molecular clone pBLV-IF2, or co-cultured with supernatants from permanently BLV-infected FLK-BLV cells or FLK-BLV cells at 4 h post transfection. CHO-K1 cells were transfected with bCAT1/pEGFP-N1 or with pEGFP-N1 as a negative control. After 48 h incubation, the cells were fixed with 3.7% formaldehyde and stained with 10 µg/mL Hoechst 33342. EGFP-expressing syncytia were evaluated and quantitated using EVOS2 fluorescence microscopy.

To evaluate whether overexpression of exogenous CAT1 would induce syncytia upon BLV infection in CHO-K1 cells, which do not normally express detectable levels CAT1 and are resistant for BLV infectious, bCAT1/pEGFP-N1 were transiently co-transfected with pBLV-IF2 into CHO-K1 cells. Interestingly, multinucleated syncytia expressing EGFP were detected in the co-transfected CHO-K1 cells, despite the fact that syncytia without EGFP expression were not observed (Fig. 2B, lower panel). In contrast, CHO-K1 cells exhibited EGFP expression with no syncytia formation when transfected with bCAT1/pEGFP-N1 alone or with negative control vector pEGFP-N1. Furthermore, BLV infection via cell-to-cell transmission using FLK-BLV cells or cell-free transmission using culture supernatants derived from FLK-BLV cells resulted in remarkable syncytia that expressed EGFP in bCAT1/pEGFP-N1-transfected CHO-K1 cells. In contrast, there was no syncytia formation when CHO-K1 cells not expressing CAT1 were infected with BLV either by cell-to-cell infection or cell-free infection or transfected with pBLV-IF2 (Fig. 2B upper panel). These results clearly indicated that bovine CAT1 expression in resistant CHO-K1 cells conferred susceptible to BLV infection.

### CAT1/SLC7A1 bound to BLV particles on the cell surface

BLV binds to cellular receptors on target cells, its membrane fusion is activated, and the virus enters the host cells. To further determine whether CAT1 acted as a cell surface receptor for BLV infection, we verified that BLV particles bound to CAT1 using flow cytometry (Fig. 3). CHO-K1 cells were transfected with bCAT1/pEGFP-N1 and incubated for 1 h or 2 h at 4°C with BLV particles purified from the culture supernatant of FLK-BLV cells. The cells were then washed and labeled with an anti-gp51 monoclonal antibody (BLV-1) followed by an allophycocyanin (APC)-conjugated anti-mouse IgG to detect the BLV particles adsorbed to the cell membrane of CHO-K1 cells. BLV-binding and EGFP-tagged CAT1 expression were analyzed using flow cytometry. The proportions of APC-EGFP double-positive cells were markedly increased in bCAT1/pEGFP-N1-transfected cells (18.7% and 23.7% at 1 h and 2 h post-incubation, respectively) compared with those of control CHO-K1 cells (0.04% and 0.24% in mock-transfected cells at 1 h and 2 h post-incubation, respectively). These results demonstrated that BLV particles bound to CAT1 at the cellular membrane of cells expressing bovine CAT1, but not to cells that did not express CAT1. These findings strongly suggest that CAT1 functioned as a BLV attachment receptor.

**Fig. 3.**
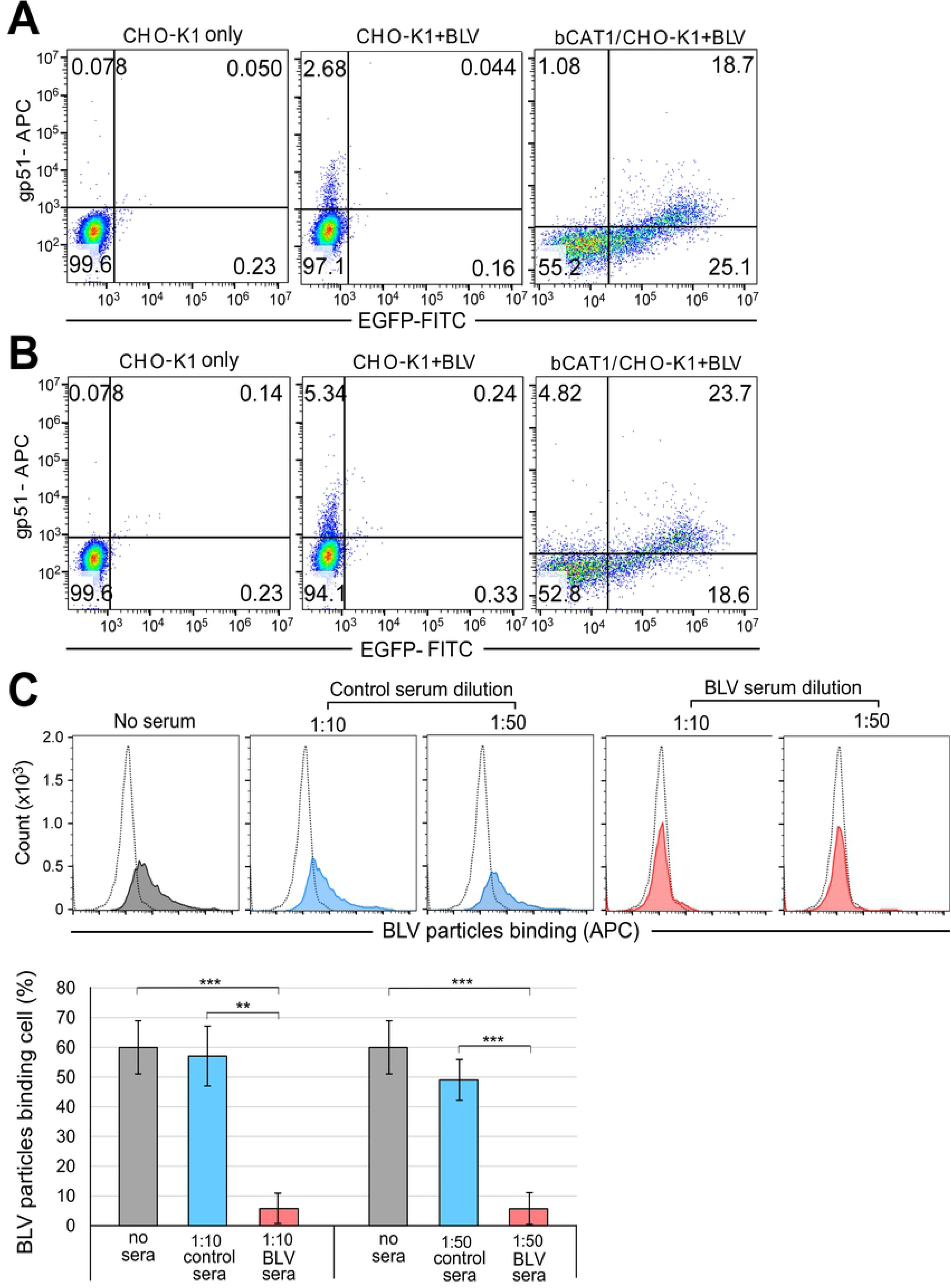
The binding ability of bovine CAT1/SLC7A1 with bovine leukemia virus (BLV) particles based on flow cytometry analysis. CHO-K1 cells were transfected with or without bCAT1/pEGFP-N1. After 48 h cultivation, the cells were incubated with BLV particles purified from the culture supernatant of FLK-BLV cells for 1 h (A) or 2 h (B) at 4 °C, and then labeled using anti-gp51 monoclonal antibody (BLV-1) followed by incubation with allophycocyanin (APC)-conjugated anti-mouse IgG. After staining, cells were analyzed using a BD Accuri™ C6 Plus flow cytometer. Each profile was separated into four quadrants, which signified single positive staining (red for BLV or green for CAT1-EGFP), double-negative staining, or double-positive staining. (C) The binding ability of CAT1/SLC7A1 and BLV particles significantly decreased by adding BLV serum. CC81-GREMG cells were added either with or without BLV serum (red) or control serum (blue) (1:10 and 1:50 dilution) and BLV particles that purified from the culture supernatant of FLK-BLV cells to bind for 2 h at 4°C. The cells were washed and then labeled using anti-gp51 antibody followed by incubation with APC-conjugated anti-mouse IgG. After staining, cells were analyzed using a BD Accuri™ C6 Plus flow cytometer. Staining cells expressed as histograms in profile. The dotted line indicated the CC81-GREMG cells only as negative control. The grey solid curve indicated cells were added BLV particles only as positive control. The percentage of BLV particles binding cells expressed as a graph (lower column). Each column and error bar represents the mean ± SD of results from triplicate experiments. The ** and *** represent *p*-values of 0.01 and 0.001, respectively (Student’s *t* test).

In addition, we used bovine serum (BLV serum) with neutralizing activity from a BLV-infected cattle [40] to examine whether the BLV serum affects the binding of CAT1-expressing CC81-GREMG cells and BLV particles. This BLV serum has been reported to inhibit syncytium formation when CC81-GREMG cells co-cultured with FLK-BLV cells [40]. CC81-GREMG cells were treated with purified BLV particles in the absence or presence of the serum for 2 h at 4°C. As shown in Fig. 3C the 60% of CC81-GREMG cells bound with BLV particles as positive control in the absence of serum (grey). When CC81-GREMG cells were treated with BLV particles in the presence of control serum (1:10 and 1:50 dilution), 57% and 49% of cells bound with BLV particles, respectively (blue). In contrast, when the cells were incubated with BLV particles in the presence of the BLV serum, proportion of BLV-bound cells was significantly reduced to about 5.8% (red). The results showed that the neutralizing serum inhibits the interaction between host cells and BLV particles.

### CAT1/SLC7A1 co-localized with BLV Env in endomembrane compartments and at the cell membrane

To further confirm the binding between BLV particles and CAT1 described above, we visualized the subcellular localization of CAT1 and Env in bCAT1/pEGFP-N1-transfected FLK-BLV cells and bCAT1/pEGFP-N1-transfected PK15/pBLV-IF2 cells using confocal laser scanning microscopy. Cells were labeled with an anti-BLV Env primary antibody followed by an Alexa Fluor 594-conjugated secondary antibody (red fluorescence) to detect cells expressing Env and stained with Hoechst 33342 (blue fluorescence) for detection of the nucleolus. EGFP-linked CAT1 was detected as green fluorescence. The intensity differences of the two signals in FLK-BLV cells and PK15/pBLV-IF2 cells were analyzed and the results depicted as a line scan (Fig. 4). In FLK-BLV cells that expressed EGFP-tagged bovine CAT1, the signal location of CAT1 (green) and Env (red) mainly coexisted in endomembrane compartments (between arrows 3 and 4 of Fig. 4a) and to a slight extent at the cell membrane (between arrows 1 and 2 of Fig. 4A). As shown in Fig. 4B, the dual labeling of CAT1 (green) and Env (red) was detected in endomembrane compartments (between arrows 2 and 3) and at the cell membrane (between arrows 1 and 2, and 4 and 5) in PK15/pBLV-IF2 cells that exogenously expressed bovine CAT1. The results showed that CAT1/SLC7A1 and Env markedly colocalized in endomembrane compartments and at the cell membrane. Taken together, these findings demonstrated a specific binding between bovine CAT1 and Env.

**Fig. 4.**
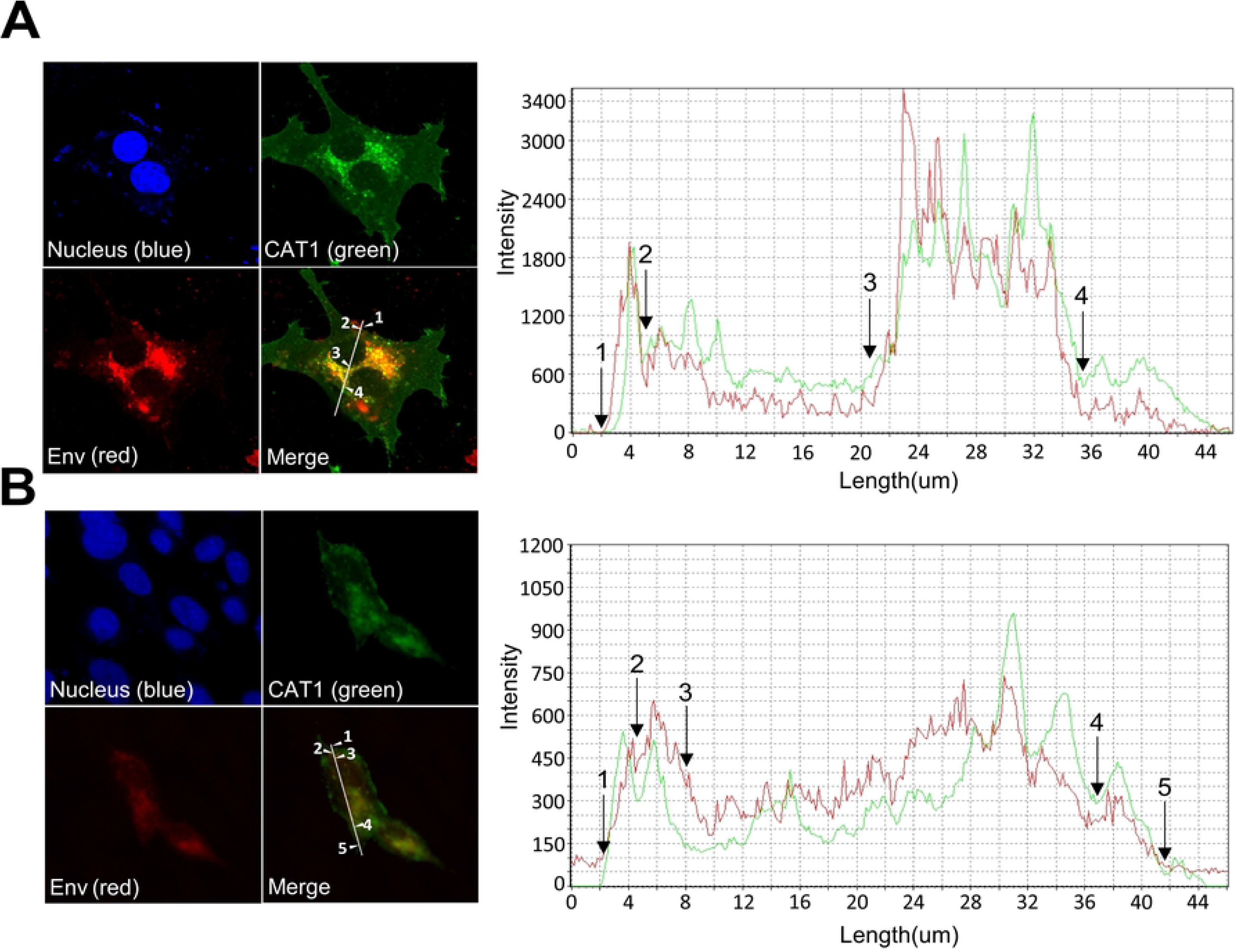
Co-localization of CAT1/SLC7A1 and bovine leukemia virus (BLV) Env protein in endomembrane compartments and at the cell membrane. Bovine CAT1/SLC7A1 expression plasmid bCAT1/pEGFP-N1 was transfected into both permanently BLV-infected FLK-BLV cells (A) and PK15/pBLV-IF2 which was stably transfected pBLV-IF2 (B), the transfected cells were grown on cover glasses and labeled with an anti-gp51 monoclonal antibody (BLV-1) followed by incubation with Alexa Fluor 594 goat anti-mouse IgG (red). Nuclei were stained with Hoechst 33342 (blue). CAT1-EGFP was visualized to determine CAT1 expression (green) in the cells. The cells were evaluated and the mean fluorescence intensities were visualized and analyzed along the line using an Olympus FV1000 laser scanning confocal fluorescence microscope. White arrows and numbers indicate that positions of endomembrane compartments and the cell membrane.

### CAT1/SLC7A1 knockdown reduced BLV infection and the binding between CAT1/SLC7A1 and BLV Env on the cell surface

To further confirm whether CAT1 acted as a cellular surface receptor for BLV infection, the impact of CAT1 knockdown on BLV infection was analyzed. To visualize the BLV infection, we used CC81-GREMG cells. CC81-GREMG cell is a CC81-derived reporter cell line harboring pBLU3_GREM_-EGFP, which contains a mutant form of the glucocorticoid response element (GRE) in the U3 region of the BLV long terminal repeat [40, 41] that specifically responses to BLV Tax expression. Knockdown with two small interfering RNAs (siRNA1 and siRNA2) that target different regions of the CAT1 mRNA was confirmed by western blot analysis 48 h post treatment using an anti-CAT1 antibody (Fig. 5A). CC81-GREMG cells in which CAT1 expression had been suppressed were inoculated with BLV using either cell-free culture supernatants derived from FLK-BLV cells (Fig. 5B upper panel) or cell-to-cell infection using FLK-BLV cells (Fig. 5B lower panel). The number of syncytia was significantly decreased in the CC81-GREMG cells treated with either CAT1-specific siRNA1 or siRNA2 compared to that in the negative control siRNA-transfected cells or in untreated cells. These results showed that knockdown of CAT1 inhibited cell-free and cell-to-cell BLV infections.

**Fig. 5.**
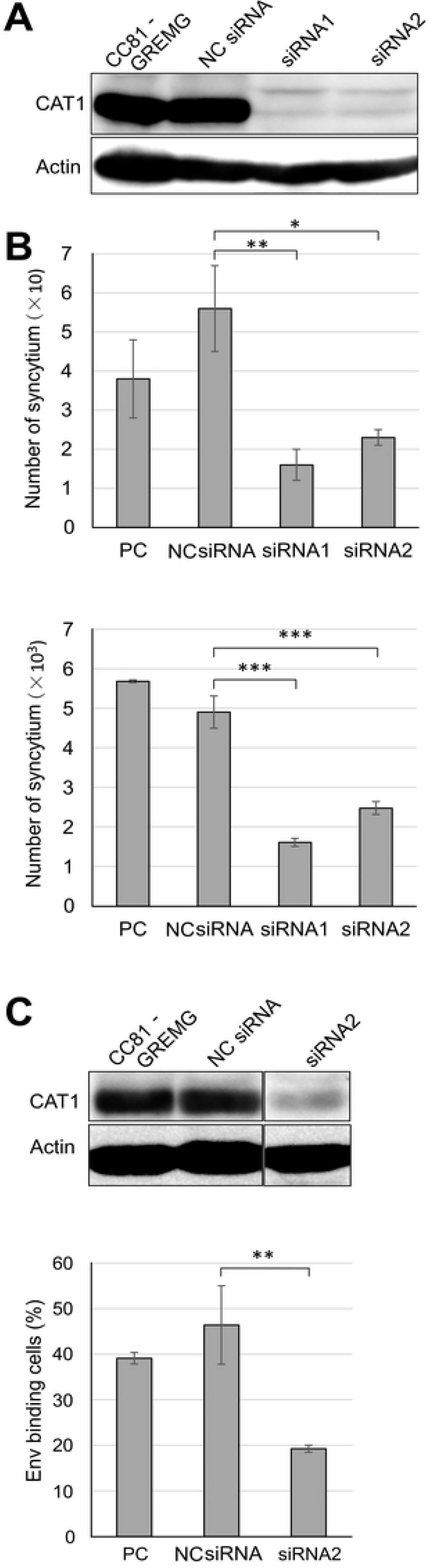
CAT1/SLC7A1 knockdown reduced bovine leukemia virus (BLV) infection. (A) CAT1/SLC7AI knockdown in CC81-GREMG cells. CC81-GREMG cells were transfected with two CAT1-specific small interfering RNAs (siRNAs) targeting different regions of CAT1/SLC7A1 or with nonspecific siRNA as a negative control. The cell extracts were prepared at 48 h post transfection and then subjected to western blot analysis using anti-CAT1 (upper panel) and anti-β-actin antibodies (lower panel). (B) CAT1/SLC7A1 knockdown suppressed cell-free and cell-to-cell BLV infection. The siRNA-transfected CC81-GREMG cells were washed and then co-cultured either with culture supernatant of FLK-BLV cells (upper) or FLK-BLV cells (lower) for 48 h and EGFP expressing syncytia were detected using EVOS2 fluorescence microscopy. Each column and error bar represents the mean ± SD of results from triplicate experiments. The *, **, and *** represent *p*-values of 0.05, 0.01, and 0.001, respectively (Student’s *t* test). (C) Env binding decreased in CAT1/SLC7A1 knockdown CC81-GREMG cells. CC81-GREMG cells were transfected with siRNA for 48 h. The transfected cells were harvested and CAT1 expression levels analyzed by western blot using an anti-CAT1 antibody. Actin was used as a loading control. CAT1 knockdown CC81-GREMG cells were added to BLV particles enriched from the culture supernatant of FLK-BLV cells and allowed to bind for 2 h at 4 °C. BLV Env binding cells were detected using anti-gp51 monoclonal antibody (BLV-1) followed by incubation with allophycocyanin (APC)-conjugated anti-mouse IgG. After staining, the cells were analyzed using a BD Accuri™ C6 Plus flow cytometry (lower panel). Each column and error bar represents the mean ± SD of results from triplicate experiments. The ** represents a *p*-value of 0.01 (Student’s *t* test).

Binding of BLV particles to the CC81-GREMG cells in which CAT1 expression had been knocked down was analyzed using the same approach as that described for Fig. 3. When CAT1 expression was reduced in CC81-GREMG cells by siRNA2 treatment compared to that in the negative control siRNA-transfected cells (Fig. 5C upper panel), the number of Env binding cells was significantly decreased (Fig. 5C lower column), indicating that the siRNA-mediated CAT1 knocked down attenuated the binding of BLV particles to target cells.

Taken together, these results indicated that CAT1 was required for BLV binding and infection. This supports the conclusion that CAT1 acted as a cell surface receptor for BLV infection.

### Species specificity of CAT1/SLC7A1 in susceptibility to BLV infection

To analyze species-specific susceptibility of CAT1 for BLV infection, CAT1 complementary DNAs (cDNAs) were generated from cell lines from eight different animal species. The cell lines included mouse NIH 3T3 cells, rat F10 cells, Chinese hamster THK cells, cat CC81 cells, sheep FLK-BLV cells, pig PK15 cells, monkey COS1 cells, and human Hela cells. EGFP-tagged CAT1 expression plasmids were constructed and CHO-K1 cells were transfected with the plasmids. The CHO-K1-transfected cells were inoculated with culture supernatants derived from FLV-BLV cells. As shown in Fig. 6A, all CAT1 proteins induced syncytia upon BLV infection. These results suggested that CAT1 did not demonstrate any species specificity for BLV infection. This was consistent with previous reports showing that BLV is able to infect cells isolated from various species *in vitro* and to propagate in various animal species *in vivo*.

**Fig. 6.**
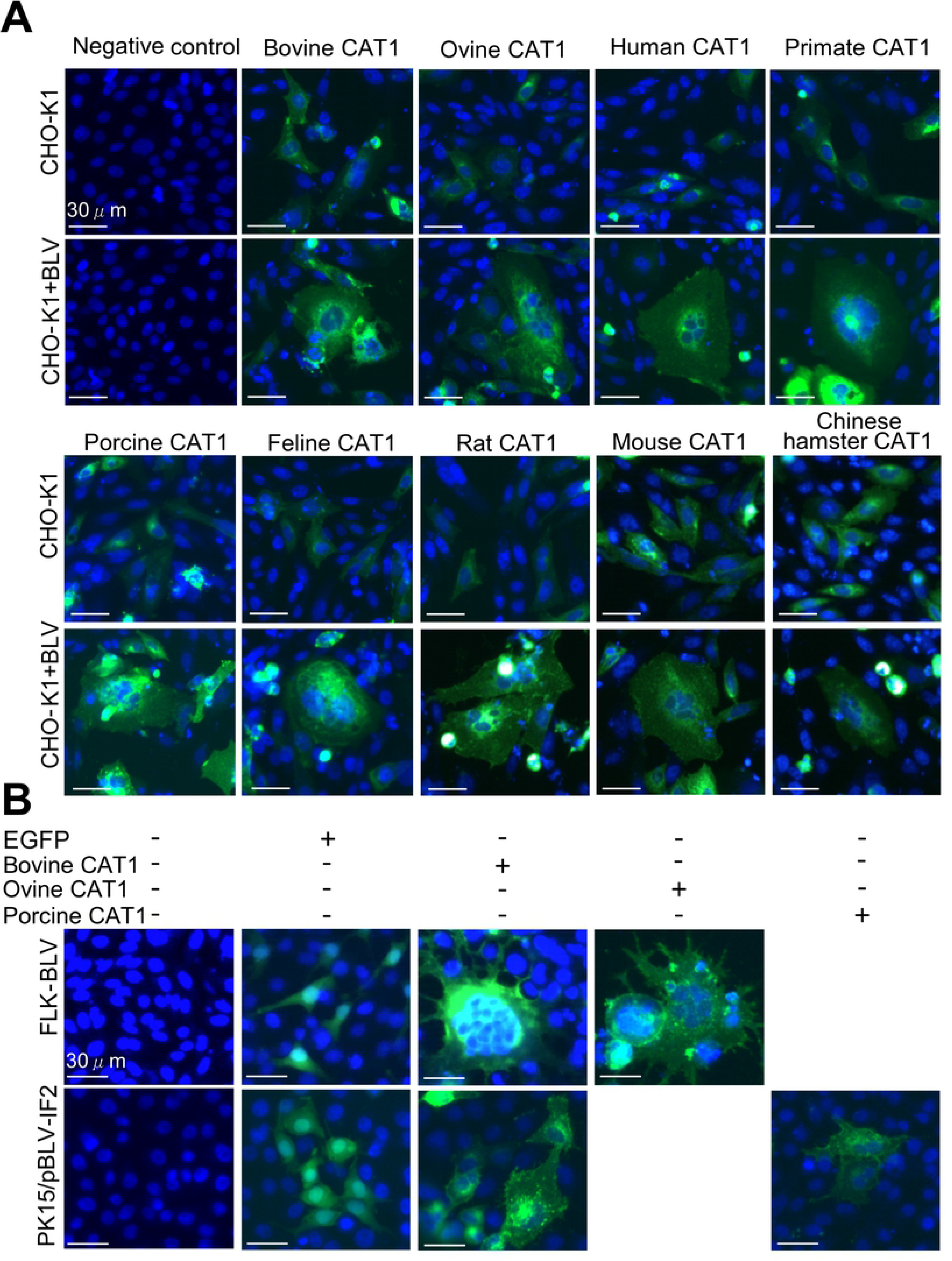
CAT1/SLC7A1 did not have species-specificity with all demonstrating susceptibility to bovine leukemia virus (BLV) infection. (A) CAT1/SLC7A1(s) from nine different animal species did not have species-specificity for BLV infection. The pEGFP-N1 expression plasmids encoding different CAT1/SLC7A1 proteins were derived using various animal species, including cattle (bovine), sheep (ovine), human, monkey (primate), pig (porcine), cat (feline), rat, mouse, and Chinese hamster. The plasmids were constructed and transfected into CHO-K1 cells. After 48 h transfection, the cells were added to culture supernatants of FLK-BLV cells and cultured for 48 h. The cells were then fixed with 3.7% formaldehyde and stained with 10 µg/mL Hoechst 33342. EGFP-expressing syncytia were detected using EVOS2 fluorescence microscopy. (B) Ovine, porcine, and bovine CAT1/SLC7A1 proteins formed syncytia in both FLK-BLV and PK15/pBLV-IF2 cells following BLV infection. The permanently BLV-infected FLK-BLV cells and PK15/pBLV-IF2 cells stably transfected pBLV-IF2 were transfected with either ovine CAT1/pEGFP-N1, porcine CAT1/pEGFP-N1, bovine CAT1/pEGFP-N1, or pEGFP-N1 and incubated for 48 h. The cells were then fixed with 3.7% formaldehyde and stained using 10 µg/mL Hoechst 33342. EGFP-expressing syncytia were detected using EVOS2 fluorescence microscopy.

To further confirm that CAT1 exhibited no species specificity for BLV infection, the ovine CAT1/pEGFP-N1 that encoded EGFP-tagged sheep CAT1 and the porcine CAT1/pEGFP-N1 that encoded EGFP-tagged pig CAT1 were transfected into FLK-BLV and PK15/pBLV-IF2 cells. Since these cells expressed BLV, it is likely that the CAT1 protein was occupied by Env protein and thereby *de novo* BLV infection did not occur. However, other mechanisms may have inhibited BLV infection. For example, host defense factors that restrict BLV infection may have been induced in the cells. As shown in Fig. 6B, syncytia containing EGFP fluorescence was observed in both CAT1-overexpressed FLK-BLV cells and CAT1-overexpressed PK15/pBLV-IF2 cells, as well as in cells transfected with bCAT1/pEGFP-N1. In contrast, syncytia formation was not detected in the pEGFP-N1-transfected FLK-BLV cells or pEGFP-N1-transfected PK15/pBLV-IF2 cells. Overexpression of CAT1 resulted in unoccupied receptor protein, resulting in the cells becoming susceptible to BLV infection. These cells did not express any inhibitors against BLV infection. These findings strongly suggested that ovine CAT1 and porcine CAT1 proteins allowed BLV infection and served as cellular receptors for BLV infection.

Taken together, these results suggested that the CAT1 proteins do not have species specificity for BLV infection, all role as a cellular receptor for BLV.

## Discussion

In the current study, we demonstrated that CAT1 also associated with BLV particles at the cell surface and functioned as a receptor for BLV infection. This is the first report demonstrating the identification of a BLV receptor. The BLVRcp1 cDNA clone corresponding to a bovine BLV receptor gene was previously isolated [42, 43]. BLVRcp1 appears to encode a protein that spans the plasma membrane once and has a molecular mass of 94 kDa. This recombinant protein, when expressed in *Escherichia coli*, binds to the BLV Env protein [42, 44] and transfection of the cDNA into mouse NIH 3T3 cells greatly increases their susceptibility to BLV infection [42]. However, the open reading frame (ORF) of the cDNA of the murine ortholog codes at a site 280 amino acids upstream of the termination codon of the ORF of the bovine ortholog [45] and the mRNA from bovine cells that hybridizes with BLVRcp1 is longer than the BLVRcp1 clone [42, 43]. In addition, mouse BLVR homolog is closely related to the δ subunit of adaptor-related protein complex AP-3 and does not associate with the cell surface [45]. Taken together, these findings suggest that this particular bovine protein is not the BLV receptor. Based on our current study, we clearly demonstrated that the CAT1 protein was the actual BLV receptor. This conclusion is based on the following findings: (i) CAT1 expression in resistant cells conferred susceptible to BLV infection, (ii) CAT1 knockdown in susceptible cells reduced BLV infection, and (iii) CAT1 exhibited specific binding to BLV Env and coexisted with BLV Env. Furthermore, GLUT1, NRP-1, and HSPG have been previously determined to be cellular receptors for HTLV, a BLV-related virus [27–29]. In addition, BLV and HTLV do not exhibit receptor interference and so it is anticipated that these viruses use distinct receptors [25]. These results also help confirm that CAT1 acted as a receptor for BLV infection.

CAT1 protein is highly regulated through transcription, mRNA stability, translation, and subcellular localization [46]. As expected, CAT1 is expressed in a wide variety of cultured cell lines since it has essential roles in basic cellular functions [46]. Indeed, CAT1 was ubiquitously expressed in cells from a variety of species and all CAT1s from the examined species were associated with susceptible to BLV infection. Therefore, the binding domain of CAT1 with BLV Env is predicted to be well conserved among various species. However, CAT1 has high diversity in the third extracellular loop near the virus-binding motif [38] for eMuLV infection. Further clarification regarding the precise roles of CAT1 and its regulation in BLV infection will help define the molecular basis of BLV entry. This includes determining the mode of binding for CAT1 with BLV Env. These results may greatly facilitate the development of therapeutic and prophylactic strategies for BLV, which has spread worldwide and is responsible for serious problems in the cattle industry.

## Materials and methods

### Cell culture and transfection

Human cervical Hela cells (RIKEN BioResource Research Center, Ibaraki, Japan), human embryonic kidney 293T cells (RIKEN BioResource Research Center, Ibaraki, Japan) feline CC81 cells (kind gift from M. Onma), the permanently BLV-infected fetal lamb kidney cell line FLK-BLV (kind gift from M. Onma) [47, 48], the calf kidney Tokushima-1 cell line CKT1(RIKEN BioResource Research Center, Ibaraki, Japan), bovine lymphoid cell line KU-1 (RIKEN BioResource Research Center, Ibaraki, Japan) [48], and African green monkey kidney fibroblast-like cell line COS1 (RIKEN BioResource Research Center, Ibaraki, Japan) were cultured in Dulbecco’s modified Eagle’s medium (DMEM; Thermo Fisher, Waltham, MA) supplemented with 10 % fetal bovine serum (FBS; Sigma-Aldrich, St. Louis, MO) at 37 °C with 5% CO_2_. The Chinese hamster ovary cell line CHO-K1 (RIKEN BioResource Research Center, Ibaraki, Japan) was cultured in Gibco™ Ham’s F-12 Nutrient Mix containing 10% FBS. The pig kidney-15 cell line PK15 (RIKEN BioResource Research Center, Ibaraki, Japan) was cultured in Gibco™ Minimum Essential Medium Eagle containing 10% FBS. For transfection of the expression plasmids and siRNAs, we used FuGENE HD (Promega, Fitchburg, WI) and Lipofectamine RNAiMAX Reagent (Invitrogen, Carlsbad, CA), according to the manufacturers’ instructions.

### Construction of CAT1/SLC7A1 expression plasmids

Total RNA was extracted from Hela, COS1, FLK-BLV, CC81, KU-1, and PK15 cells using TRIzol® LS Reagent (Thermo Fisher Scientific, Carlsbad, CA) and reverse-transcribed into cDNA using a High-Capacity RNA-to-cDNA Kit (Thermo Fisher Scientific, Vilnius, Lithuania). The SLC7A1 cDNA from each sample was amplified by polymerase chain reaction (PCR) using PrimeSTAR GXL DNA polymerase (Takara Bio, Otsu, Japan). The amplification profile included 98 °C for 2 min followed by 30 cycles of 98 °C for 15 s, 55 °C for 5–15 s, and 72 °C for 30 s. The primer sets used are shown in supplementary Table 1. Each PCR amplified product of SLC7A1 was purified by using a FastGene Gel/PCR Extraction Kit (Nippon Genetics, Tokyo, Japan) and cloned into a pEGFP-N1 vector that encodes EGFP [49] using an In-Fusion HD Cloning Kit (TaKaRa, Mountain View, CA). The mouse, rat, and Chinese hamster SLC7A1s were digested from their respective expression plasmids SLC7A1/pTarkeT [50] and ligated into pEGFP-N1. All DNA sequences of the constructed expression plasmids were confirmed by sequencing using the BigDye Terminator v.3.1 Cycle Sequencing Kit (Applied Biosystems, Waltham, MA) and analyzed using GENETYX ver. 10 software (Genetyx, Tokyo, Japan) and compared to GenBank.

### Western blotting

Cell lysates of Hela, 293T, CC81, CKT1, KU-1, FLK-BLV, PK15, and CHO-K1 cultures were extracted on ice for 1 h with radioimmunoprecipitation assay (RIPA) buffer containing 50 mM Tris-HCl/pH 7.5, 150 mM NaCl, 1% TritonX-100, 0.1% sodium dodecyl sulphate (SDS; Sigma-Aldrich, Saint Louis, MO), 1% sodium deoxycholate (Sigma-Aldrich Mannheim, Germany), and 1% protease inhibitor (Sigma-Aldrich). The protein concentrations of the lysates were then quantified using a bicinchoninic acid protein kit (Thermo Fisher Scientific, Rockford, IL). Aliquots containing 12 µg of total cellular lysates for each sample were denatured at 100 °C for 5 min and separated using 8% sodium dodecyl sulfate polyacrylamide gel electrophoresis (SDS-PAGE). Samples were transferred to a membrane, blocked with 5% skim milk in phosphate-buffered saline (PBS)-T buffer, and then incubated with a rabbit anti-CAT1 antibody (Abcam, Cambridge, MA). After labeling with the primary antibody, the blots were incubated with a secondary horseradish peroxidase (HRP)-conjugated goat anti-rabbit IgG (Amersham Biosciences, Piscataway, NJ).

### Syncytium formation assay

Cells which used for syncytium formation assays were prepared as follows: (`Ⅰ`) CHO-K1 cells (5 × 10^4^) were transfected with bovine CAT1/pEGFP-N1 (bCAT1/pEGFP-N1), human CAT1/pEGFP-N1, primate CAT1/pEGFP-N1, ovine CAT1/pEGFP-N1, porcine CAT1/pEGFP-N1, feline CAT1/pEGFP-N1, mouse CAT1/pEGFP-N1, rat CAT1/pEGFP-N1, or Chinese hamster CAT1/pEGFP-N1 and then added culture supernatants of FLK-BLV cells. (`Ⅱ`) The CHO-K1 cells (5 × 10^4^) were co-transfected with infectious molecular clone pBLV-IF2 [17, 51] and bCAT1/pEGFP-N1 expression plasmid. (`Ⅲ`) FLK-BLV cells and PK15/pBLV-IF2 cells were transfected with bCAT1/pEGFP-N1 expression plasmid. (`Ⅳ`) FLK-BLV cells and PK15/pBLV-IF2 cells were transfected with ovine CAT1/pEGFP-N1 and porcine CAT1/pEGFP-N1 expression plasmid, respectively. After 48 h of incubation post transfection, all cells were washed with PBS and fixed with PBS containing 3.7% formaldehyde and 10 µg/mL Hoechst 33342 (ImmunoChemistry Technologies LLC, Bloomington, MN). The fixed cells were observed for fluorescent-positive syncytia using an EVOS FL Auto 2 Cell Imaging System (Thermo Fisher Scientific).

CC81-GREMG cells [40, 41] were transfected with anti-CAT1/SLC7A1 siRNAs and then co-cultured with the culture supernatant of FLK-BLV cells or FLK-BLV cells. The number of syncytia with EGFP expression was counted as described above.

### Immunofluorescence assay

FLK-BLV and PK15/pBLV-IF2 cells (5 × 10^4^) were seeded onto cover glasses in 12-well plates and transfected with 2 µg of bCAT1/pEGFP-N1 expression plasmid. After 48 h transfection, the cover glasses were removed for immunofluorescence staining. The staining included the following steps: the cells were fixed with 4% paraformaldehyde, treated by 0.1% Triton-X, and blocked with 5% skim milk. For Env protein detection, the cells were incubated with a primary anti-Env monoclonal antibody (Mab, BLV-1; VMRD, Pullman, WA) followed by incubation with Alexa Fluor 594 goat anti-mouse IgG (Invitrogen). Nuclei were stained with Hoechst 33342 in the dark. The mean fluorescence intensities of Alexa Fluor 594 and EGFP in the cells were visualized and analyzed along the line using an FV1000 confocal laser scanning microscope (Olympus, Tokyo, Japan).

### Analysis of CAT1/SLC7A1 and BLV particle binding

BLV particles were purified from the culture supernatant of FLK-BLV cells using ultracentrifugation and re-suspended in F-12 medium containing 4 µg/ml of polybrene solution (Sigma-Aldrich). CHO-K1 cells were seeded 2 × 10^5^ cells/well into 6-well plate one day prior to use and then transfected with 4 µg of bCAT1/pEGFP-N1 expression plasmid. At 48 h post transfection, the cells were incubated with 200 µl aliquots of the BLV particles at 4 °C. After washing five times with ice-cold fresh culture medium to remove unbound virus particles, the cells were treated with 2 mM EDTA/PBS and harvested using 1 ml 0.5% FBS in PBS. To determine the binding of CHO-K1/CAT1 with BLV-Env, harvested cells were incubated on ice with an anti-Env Mab (BLV-1) for 1 h or 2 h and then subsequently incubated for 30 min on ice with an APC-conjugated rat anti-mouse IgG (BD Biosciences, CA). Stained cells were analyzed using a BD Accuri^TM^ C6 Plus with a Sampler flow cytometer (BD Biosciences, San Jose, California). The data were analyzed using FlowJo Single Cell Analysis Software v10 (FlowJo, LLC, Ashland, OH). As a negative control, bCAT1/pEGFP-N1 transfected cells were incubated with secondary antibody only.

### Neutralization assay

BLV serum was collected from BLV-infected cattle with lymphoma, and control serum was isolated from BLV-negative healthy cattle. The serum were 10- and 50-fold diluted with fresh culture medium. CC81-GREMG cells (2 x 10^5^) were incubated with BLV particles in the absence or presence of the serum for 2 h at 4 °C. BLV particle-bound cells were detected as described above.

### CAT1/SLC7A1 knockdown assay

Silencer select siRNAs (Invitrogen) targeting the CAT1/SLC7A1 gene were designed by GeneAssist™ Custom siRNA Builder (Thermo Fisher Scientific) and synthesized. Silencer select siRNA Negative Control Med GC (Invitrogen) was used as a negative control siRNA. A mixture of siRNAs was transfected into CC81-GREMG cells to knock down CAT1/SLC7A1 expression. After 48 h transfection, the CC81-GREMG cells were cocultured with BLV particles and binding between CAT1 and BLV Env was assayed as described above. The transfected CC81-GREMG cells were treated with 2 mM EDTA/PBS and harvested for western blot analysis to quantify CAT1/SLC7A1 levels using an anti-CAT1 antibody.

## Acknowledgments

We thank the members of Viral Infectious Disease Unit, RIKEN for technical assistance and helpful suggestions, and the Support Unit for Biomaterial Analysis, RIKEN BSI Research Resources Center for help with sequence analysis.

This work was supported by Grants-in-Aid for Scientific Research [A (16H02590)] from the Japan Society for the Promotion of Science (JSPS), by a grant from the Project of the NARO Bio-oriented Technology Research Advancement Institution (the special scheme project on regional developing strategy; Grant No. 16817983), by a grant from the Project of the NARO Bio-oriented Technology Research Advancement Institution (the special scheme project on vitalizing management entities of agriculture, forestry and fisheries; Grant No. 16930548), and by the Japan Society for the JSPS Postdoctoral Fellowship (Grant No. 16F16404).

The funders had no role in study design, data collection and analysis, decision to publish, or preparation of the manuscript.

## Supporting information Legends

S1 Table. Oligonucleotide primers used for construction of nine species of CAT1/SLC7A1 expression plasmids and two kinds of CAT1-specific small interfering RNA (siRNA).

## Author Contributions

L.B. participated in the execution of all experiments, analyzed the data, and drafted the manuscript. H.S. participated in construction of the CAT1/SLC7A1 expression plasmids. S.W. participated in analyzing the data. K.Y. participated in construction of the CAT1/SLC7A1 expression plasmids and drafted the manuscript. Y.A. participated in the execution of all experiments, participated in experimental design, coordinated the experiments, and drafted the manuscript. All authors reviewed and approved the final manuscript.

## Conflicts of interest

The authors declare no conflicts of interest.

## References

1. Aida Y, Murakami H, Takahashi M, Takeshima SN. Mechanisms of pathogenesis induced by bovine leukemia virus as a model for human T-cell leukemia virus. Front Microbiol. 2013;4:328. doi: 10.3389/fmicb.2013.00328. PubMed PMID: 24265629; PubMed Central PMCID: PMCPMC3820957.

2. APHIS. Bovine Leukosis Virus (BLV) on U.S. Dairy Operations, 2007. United states department of agriculture. 2008.

3. Miller JM, Miller LD, Olson C, Gillette KG. Virus-like particles in phytohemagglutinin-stimulated lymphocyte cultures with reference to bovine lymphosarcoma. J Natl Cancer Inst. 1969;43(6):1297–305. PubMed PMID: 5408805.

4. Gillet N, Florins A, Boxus M, Burteau C, Nigro A, Vandermeers F, et al. Mechanisms of leukemogenesis induced by bovine leukemia virus: prospects for novel anti-retroviral therapies in human. Retrovirology. 2007;4:18. doi: 10.1186/1742-4690-4-18. PubMed PMID: 17362524; PubMed Central PMCID: PMCPMC1839114.

5. Polat M, Takeshima SN, Aida Y. Epidemiology and genetic diversity of bovine leukemia virus. Virol J. 2017;14(1):209. doi: 10.1186/s12985-017-0876-4. PubMed PMID: 29096657; PubMed Central PMCID: PMCPMC5669023.

6. Aida Y, Miyasaka M, Okada K, Onuma M, Kogure S, Suzuki M, et al. Further phenotypic characterization of target cells for bovine leukemia virus experimental infection in sheep. Am J Vet Res. 1989;50(11):1946–51. PubMed PMID: 2559634.

7. Nagaoka Y, Kabeya H, Onuma M, Kasai N, Okada K, Aida Y. Ovine MHC class II DRB1 alleles associated with resistance or susceptibility to development of bovine leukemia virus-induced ovine lymphoma. Cancer Res. 1999;59(4):975–81. PubMed PMID: 10029093.

8. Olson C, Kettmann R, Burny A, Kaja R. Goat lymphosarcoma from bovine leukemia virus. J Natl Cancer Inst. 1981;67(3):671–5. PubMed PMID: 6268881.

9. Mammerickx M, Portetelle D, Burny A. Experimental cross-transmissions of bovine leukemia virus (BLV) between several animal species. Zentralbl Veterinarmed B. 1981;28(1):69–81. PubMed PMID: 6263020.

10. Altanerova V, Portetelle D, Kettmann R, Altaner C. Infection of rats with bovine leukaemia virus: establishment of a virus-producing rat cell line. J Gen Virol. 1989;70 (Pt 7):1929–32. doi: 10.1099/0022-1317-70-7-1929. PubMed PMID: 2544673.

11. Altanerova V, Ban J, Altaner C. Induction of immune deficiency syndrome in rabbits by bovine leukaemia virus. AIDS. 1989;3(11):755–8. PubMed PMID: 2559751.

12. Altanerova V, Ban J, Kettmann R, Altaner C. Induction of leukemia in chicken by bovine leukemia virus due to insertional mutagenesis. Arch Geschwulstforsch. 1990;60(2):89–96. PubMed PMID: 2160229.

13. Buehring GC, Philpott SM, Choi KY. Humans have antibodies reactive with Bovine leukemia virus. AIDS Res Hum Retroviruses. 2003;19(12):1105–13. doi: 10.1089/088922203771881202. PubMed PMID: 14709247.

14. Buehring GC, Shen HM, Jensen HM, Choi KY, Sun D, Nuovo G. Bovine leukemia virus DNA in human breast tissue. Emerg Infect Dis. 2014;20(5):772–82. doi: 10.3201/eid2005.131298. PubMed PMID: 24750974; PubMed Central PMCID: PMCPMC4012802.

15. Buehring GC, Shen H, Schwartz DA, Lawson JS. Bovine leukemia virus linked to breast cancer in Australian women and identified before breast cancer development. PLoS One. 2017;12(6):e0179367. doi: 10.1371/journal.pone.0179367. PubMed PMID: 28640828; PubMed Central PMCID: PMCPMC5480893.

16. Buehring GC, Shen HM, Jensen HM, Jin DL, Hudes M, Block G. Exposure to Bovine Leukemia Virus Is Associated with Breast Cancer: A Case-Control Study. PLoS One. 2015;10(9):e0134304. doi: 10.1371/journal.pone.0134304. PubMed PMID: 26332838; PubMed Central PMCID: PMCPMC4557937.

17. Inabe K, Ikuta K, Aida Y. Transmission and propagation in cell culture of virus produced by cells transfected with an infectious molecular clone of bovine leukemia virus. Virology. 1998;245(1):53–64. doi: 10.1006/viro.1998.9140. PubMed PMID: 9614867.

18. Webster RG, Granoff A. Encyclopedia of Virology: A-Fur: Academic Press; 1994.

19. Sagata N, Yasunaga T, Ohishi K, Tsuzuku-Kawamura J, Onuma M, Ikawa Y. Comparison of the entire genomes of bovine leukemia virus and human T-cell leukemia virus and characterization of their unidentified open reading frames. EMBO J. 1984;3(13):3231–7. PubMed PMID: 6098469; PubMed Central PMCID: PMCPMC557842.

20. Johnston ER, Radke K. The SU and TM envelope protein subunits of bovine leukemia virus are linked by disulfide bonds, both in cells and in virions. J Virol. 2000;74(6):2930–5. PubMed PMID: 10684314; PubMed Central PMCID: PMCPMC111788.

21. Hunter E. Viral Entry and Receptors. In: Coffin JM, Hughes SH, Varmus HE, editors. Retroviruses. Cold Spring Harbor (NY)1997.

22. Callebaut I, Voneche V, Mager A, Fumiere O, Krchnak V, Merza M, et al. Mapping of B-neutralizing and T-helper cell epitopes on the bovine leukemia virus external glycoprotein gp51. J Virol. 1993;67(9):5321–7. PubMed PMID: 7688821; PubMed Central PMCID: PMCPMC237931.

23. Bruck C, Mathot S, Portetelle D, Berte C, Franssen JD, Herion P, et al. Monoclonal antibodies define eight independent antigenic regions on the bovine leukemia virus (BLV) envelope glycoprotein gp51. Virology. 1982;122(2):342–52. PubMed PMID: 6183819.

24. Weiss RA, Tailor CS. Retrovirus receptors. Cell. 1995;82(4):531–3. PubMed PMID: 7664331.

25. Sommerfelt MA. Retrovirus receptors. J Gen Virol. 1999;80 (Pt 12):3049–64. doi: 10.1099/0022-1317-80-12-3049. PubMed PMID: 10567635.

26. Manel N, Kim FJ, Kinet S, Taylor N, Sitbon M, Battini JL. The ubiquitous glucose transporter GLUT-1 is a receptor for HTLV. Cell. 2003;115(4):449–59. PubMed PMID: 14622599.

27. Ghezzi PC, Dolcini GL, Gutierrez SE, Bani PC, Torres JO, Arroyo GH, et al. [Bovine leukemia virus (BLV): prevalence in the Cuenca Lechera Mar y Sierras from 1994 to 1995]. Rev Argent Microbiol. 1997;29(3):137–46. PubMed PMID: 9411488.

28. Jones KS, Petrow-Sadowski C, Bertolette DC, Huang Y, Ruscetti FW. Heparan sulfate proteoglycans mediate attachment and entry of human T-cell leukemia virus type 1 virions into CD4+ T cells. J Virol. 2005;79(20):12692–702. doi: 10.1128/JVI.79.20.12692-12702.2005. PubMed PMID: 16188972; PubMed Central PMCID: PMCPMC1235841.

29. Ghez D, Lepelletier Y, Jones KS, Pique C, Hermine O. Current concepts regarding the HTLV-1 receptor complex. Retrovirology. 2010;7:99. doi: 10.1186/1742-4690-7-99. PubMed PMID: 21114861; PubMed Central PMCID: PMCPMC3001707.

30. Jones KS, Lambert S, Bouttier M, Benit L, Ruscetti FW, Hermine O, et al. Molecular aspects of HTLV-1 entry: functional domains of the HTLV-1 surface subunit (SU) and their relationships to the entry receptors. Viruses. 2011;3(6):794–810. doi: 10.3390/v3060794. PubMed PMID: 21994754; PubMed Central PMCID: PMCPMC3185769.

31. Pinon JD, Klasse PJ, Jassal SR, Welson S, Weber J, Brighty DW, et al. Human T-cell leukemia virus type 1 envelope glycoprotein gp46 interacts with cell surface heparan sulfate proteoglycans. J Virol. 2003;77(18):9922–30. PubMed PMID: 12941902; PubMed Central PMCID: PMCPMC224595.

32. Mueckler M. Facilitative glucose transporters. Eur J Biochem. 1994;219(3):713–25. PubMed PMID: 8112322.

33. Albritton LM, Tseng L, Scadden D, Cunningham JM. A putative murine ecotropic retrovirus receptor gene encodes a multiple membrane-spanning protein and confers susceptibility to virus infection. Cell. 1989;57(4):659–66. PubMed PMID: 2541919.

34. Kim JW, Closs EI, Albritton LM, Cunningham JM. Transport of cationic amino acids by the mouse ecotropic retrovirus receptor. Nature. 1991;352(6337):725–8. doi: 10.1038/352725a0. PubMed PMID: 1652100.

35. Wang H, Kavanaugh MP, North RA, Kabat D. Cell-surface receptor for ecotropic murine retroviruses is a basic amino-acid transporter. Nature. 1991;352(6337):729–31. doi: 10.1038/352729a0. PubMed PMID: 1908564.

36. Albritton LM, Kim JW, Tseng L, Cunningham JM. Envelope-binding domain in the cationic amino acid transporter determines the host range of ecotropic murine retroviruses. J Virol. 1993;67(4):2091–6. PubMed PMID: 8445722; PubMed Central PMCID: PMCPMC240296.

37. Yoshimoto T, Yoshimoto E, Meruelo D. Identification of amino acid residues critical for infection with ecotropic murine leukemia retrovirus. J Virol. 1993;67(3):1310–4. PubMed PMID: 8382297; PubMed Central PMCID: PMCPMC237498.

38. Kakoki K, Shinohara A, Izumida M, Koizumi Y, Honda E, Kato G, et al. Susceptibility of muridae cell lines to ecotropic murine leukemia virus and the cationic amino acid transporter 1 viral receptor sequences: implications for evolution of the viral receptor. Virus Genes. 2014;48(3):448–56. doi: 10.1007/s11262-014-1036-1. PubMed PMID: 24469466.

39. Bae EH, Park SH, Jung YT. Role of a third extracellular domain of an ecotropic receptor in Moloney murine leukemia virus infection. J Microbiol. 2006;44(4):447–52. PubMed PMID: 16953181.

40. Sato H, Watanuki S, Bai L, Borjigin L, Ishizaki H, Matsumoto Y, et al. A sensitive luminescence syncytium induction assay (LuSIA) based on a reporter plasmid containing a mutation in the glucocorticoid response element in the long terminal repeat U3 region of bovine leukemia virus. Virol J. 2019;16(1):66. doi: 10.1186/s12985-019-1172-2. PubMed PMID: 31109347.

41. Sato H, Watanuki S, Murakami H, Sato R, Ishizaki H, Aida Y. Development of a luminescence syncytium induction assay (LuSIA) for easily detecting and quantitatively measuring bovine leukemia virus infection. Arch Virol. 2018;163(6):1519–30. doi: 10.1007/s00705-018-3744-7. PubMed PMID: 29455325.

42. Ban J, Portetelle D, Altaner C, Horion B, Milan D, Krchnak V, et al. Isolation and characterization of a 2.3-kilobase-pair cDNA fragment encoding the binding domain of the bovine leukemia virus cell receptor. J Virol. 1993;67(2):1050–7. PubMed PMID: 8380453; PubMed Central PMCID: PMCPMC237460.

43. Ban J, Truong AT, Horion B, Altaner C, Burny A, Portetelle D, et al. Isolation of the missing 5’-end of the encoding region of the bovine leukemia virus cell receptor gene. Arch Virol. 1994;138(3-4):379–83. PubMed PMID: 7998843.

44. Orlik O, Ban J, Hlavaty J, Altaner C, Kettmann R, Portetelle D, et al. Polyclonal bovine sera but not virus-neutralizing monoclonal antibodies block bovine leukemia virus (BLV) gp51 binding to recombinant BLV receptor BLVRcp1. J Virol. 1997;71(4):3263–7. PubMed PMID: 9060692; PubMed Central PMCID: PMCPMC191461.

45. Suzuki T, Ikeda H. The mouse homolog of the bovine leukemia virus receptor is closely related to the delta subunit of adaptor-related protein complex AP-3, not associated with the cell surface. J Virol. 1998;72(1):593–9. PubMed PMID: 9420263; PubMed Central PMCID: PMCPMC109412.

46. Closs EI, Boissel JP, Habermeier A, Rotmann A. Structure and function of cationic amino acid transporters (CATs). J Membr Biol. 2006;213(2):67–77. doi: 10.1007/s00232-006-0875-7. PubMed PMID: 17417706.

47. Van Der Maaten MJ, Miller JM. Replication of bovine leukemia virus in monolayer cell cultures. Bibl Haematol. 1975;(43):360–2. PubMed PMID: 183699.

48. Onuma M, Koyama H, Aida Y, Okada K, Ogawa Y, Kirisawa R, et al. Establishment of B-cell lines from tumor of enzootic bovine leukosis. Leuk Res. 1986;10(6):689–95. PubMed PMID: 3012214.

49. Cormack BP, Valdivia RH, Falkow S. FACS-optimized mutants of the green fluorescent protein (GFP). Gene. 1996;173(1 Spec No):33–8. PubMed PMID: 8707053.

50. Kubo Y, Ono T, Ogura M, Ishimoto A, Amanuma H. A glycosylation-defective variant of the ecotropic murine retrovirus receptor is expressed in rat XC cells. Virology. 2002;303(2):338–44. PubMed PMID: 12490395.

51. Inabe K, Nishizawa M, Tajima S, Ikuta K, Aida Y. The YXXL sequences of a transmembrane protein of bovine leukemia virus are required for viral entry and incorporation of viral envelope protein into virions. J Virol. 1999;73(2):1293–301. PubMed PMID: 9882334; PubMed Central PMCID: PMCPMC103953.

